# RNA sequencing-based cell proliferation analysis across 19 cancers identifies a subset of proliferation-informative cancers with a common survival signature

**DOI:** 10.1101/063057

**Authors:** Ryne C. Ramaker, Brittany N. Lasseigne, Andrew A. Hardigan, Laura Palacio, David S. Gunther, Marie K. Kirby, Richard M. Myers, Sara J. Cooper

## Abstract

Despite advances in cancer diagnosis and treatment strategies, robust prognostic signatures remain elusive in most cancers. Cell proliferation has long been recognized as a prognostic marker in cancer, but it has not been thoroughly investigated across multiple cancers. Here we explore the role of cell proliferation across 19 cancers (n=6,581 patients) using tissue-based RNA sequencing from The Cancer Genome Atlas project by employing a ‘proliferative index’ derived from gene expression associated with *PCNA* expression. This proliferative index is significantly associated with patient survival (Cox, p-value<0.05) in 7/19 cancers, which we have defined as ‘proliferation-informative cancers’ (PICs). In PICs the proliferative index is strongly correlated with tumor stage and nodal invasion. PICs paradoxically demonstrate reduced baseline expression of proliferation machinery relative to non-PICs suggesting that non-PICs saturate their proliferative capacity early in tumor development and allow other factors to dictate prognostic outcomes. We also identify chemotherapies whose efficacy is correlated with proliferation index and highlight drugs capable of inhibiting proliferation associated expression. Additionally, we find that proliferative index is significantly associated with gross somatic mutation burden (Spearman, p=1.76×10^−23^) as well mutations in individual driver genes. This analysis provides a comprehensive characterization of tumor proliferation rates and their association with disease progression and prognosis across cancer types and highlights specific cancers that may be particularly susceptible to improved targeting of this classic cancer hallmark.

## Introduction

A fundamental characteristic of cancer cells is their ability to maintain the capacity to proliferate, bypassing the homeostatic signaling network controlling cell division in normal tissue. The capacity to “sustain proliferative signaling,” “enable replicative immortality,” and “evade growth suppressors” represent 3 of the original 6 hallmarks of cancer, and histological techniques examining the number of mitotic cells present in tumor biopsies have been used clinically to assess tumor grade for several decades [1,2]. Although proliferation is a clear hallmark of cancer, tumor evolutionary tradeoffs may exist in certain tumor types or stages that prioritize resources for other survival phenotypes like metastasis [3,4], angiogenesis [5–7], immune system evasion [8,9], drug efflux [10,11], DNA repair [12,13], drug resistance [14], or reactive oxygen species (ROS) regulation [15]. Characterizing these tradeoffs is critical for a complete understanding tumor progression and the development of appropriate therapies [16].

Early studies comparing tumor with adjacent normal tissue identified expression changes in genes controlling cell proliferation as some of the largest and most consistent cancer alterations and further associated proliferation signatures with poor patient prognosis and advanced tumor grade[17–22]. Recent large-scale sequencing efforts have described driver mutations that hijack normal proliferation machinery. For example, approximately 40% of melanomas possess activating *BRAF* mutations that modulate proliferation by constitutively activating the downstream mitogen activated protein kinase (*MAPK*) pathway[23]. Multiple tumor types also harbor activating mutations in phosphoinositide 3-kinase (*PI3K*) that hyperactivate *AKT/mTOR* signaling and several other pathways important for regulating proliferation[24]. Accordingly, a majority of cytotoxic chemotherapies seek to preferentially target the increased proliferation rate of cancer cells by damaging DNA in dividing cells or impairing vital replication machinery[25,26].

Venet, et al. recently derived a general marker of proliferation, ‘metaPCNA’, by identifying the top 1% of genes most positively correlated with the proliferation marker *PCNA* across 36 tissue types and demonstrated that it significantly outperformed a majority of prognostic signatures developed for breast cancer[27,28]. Further highlighting the importance of proliferation rate, the authors determined a majority of variation in breast cancer transcriptomes is correlated with proliferation and most random gene sets are significantly associated with breast cancer outcome due to their inherent relationship with a broad underlying proliferation signature [27,28]. In this study we examine the relative importance of proliferation to disease progression and patient prognosis across cancers using RNA-sequencing (RNA-seq) profiles from 19 cancers in 6,581 patients catalogued in The Cancer Genome Atlas (TCGA). We contrast these with 30 normal tissues from 8,553 patients from the Genotype-Tissue Expression (GTEx) project to investigate proliferation rates across tissues types and disease stages. We also describe chemotherapies whose efficacy is associated with cellular proliferation rates and highlight drugs that appear significantly alter proliferation at the transcription level. We demonstrate a strong relationship between tumor proliferation signatures and somatic mutation burden and identify single nucleotide variants associated with proliferative phenotype across cancers. The relative prognostic power of the metaPCNA proliferation index (PI), common clinical annotations, optimized survival models, and random transcript sets are compared within and across each cancer. Finally, we provide on open-source R package for future studies which calculates and analyzes PI across a user’s dataset and compares a user’s model with PI.

## Methods

### TCGA and GTEx Data acquisition

RNA-seq and associated patient clinical data was obtained from the TCGA data portal (tcga-data.nci.nih.gov) in June 2015. (Supplemental Table 1) Level 3 RNASeqV2 raw count data was used for downstream analysis. Relevant clinical information for each patient was obtained from the associated “clinical_patient” and “clinical_follow_up” files, with survival time calculated as the maximum “days_to_death” or “days_to_last_followup” column value from the “clinical_patient” file or any “clinical_follow_up” file. All staging information was obtained from the “pathologic_T”, “pathologic_N”, and “pathologic_M” columns in the “clinical_patient” file. GTEx(gtexportal.org) V6 RNA-seq data for all available tissues was obtained in January 2016 (Supplemental Table 2).

**All analysis was performed using R** [29] **(Version 3.2.1) with RStudio** [30] **(Version 0.99.891).**

### Data normalization and PI calculation

The PI was calculated as previously described by Venet et. al. Briefly, a sample’s PI was defined as the median expression value of the original 131 genes found to be most associated with *PCNA* expression across 36 tissue types. For cross-cancer or crosstissue comparisons, raw count reads were normalized to counts-per-million (CPM) prior to PI calculation. For intra-cancer analyses, raw counts were variance stabilized using the ‘DESeq2’[31] (Version 1.8.2) package function “varianceStabilizingTransformation” prior to PI calculation or survival analysis

### PI comparisons and survival association analysis

All cross-sample PI comparisons were conducted with two-sided wilcox tests via the base ‘stats’[29] (version 3.2.1) package wilcox.test function. PI-survival associations were determined using survival[32,33] (version 2.38-3) and survcomp[34,35] (version 1. 18.0) packages. Cox regressions were performed with the coxph function to regress overall patient survival on PI and Wald test p-values were reported. Kaplan-Meier curves were generated for tumors in the top and bottom quartiles of PI using the survfit function and significant differences between survival curves were assessed with the survdiff function. Dendrograms of cancer clustering based on negative log10 Cox regression p-values were constructed with the hclust function using Ward clustering. A heatmap of cross-cancer survival associated genes (uncorrected p-value<0.05 for at least 9/19 cancers) was generated on negative log10 Cox regression p-values generated for each transcript measured in TCGA Level 3 data. Models that failed to converge were assigned a p-value of 1. The heatmap was generated with the R gplots[36] (version 2.17.0) heatmap.2 function using Euclidean distance measurement and Ward clustering.

### Pathway analysis

Pathway analysis was conducted on the 162 cross-cancer survival associated genes with uncorrected Cox p-values<0.05 across all PICs using the Database for Annotation, Visualization and Integrated Discovery (DAVID, v6.7)[37,38] pathway analysis with default settings. All unique gene names available in the TCGA Level 3 count data were used as a background for analysis. Gene ontology enrichment analysis of expression-survival associations in each cancer was conducted with GOrilla (http://cbl-gorilla.cs.technion.ac.il) in “single ranked list of genes” mode. GO terms were condensed into broader categories for visualization with REVIGO (http://revigo.irb.hr).

### Cross-cancer survival model

Variance stabilized transcript count data was scaled within each cancer prior to combining cohorts for all cross-cancer survival model generation. For each cancer, the 18 shortest surviving patients who succumbed to disease and the 18 longest surviving patients were identified for initial analyses. Only 18 patients were selected because this represented the top and bottom quartiles of the mesothelioma cohort, the smallest cohort included in this study. Patients were indexed as “1” if they were in the shortest/longest overall survival analysis and “0” if they were not. We then generated two models: the first included all 19 cancers and the second included only PIC cancers (KIRC, ACC, LGG, KIRP, MESO, PAAD, and LUAD). PICs were defined as cancers with Bonferroni-corrected PI Cox regression p-value of less than 0.05 and also as the cancers who clustered together when considering only the cross-cancer significant survival transcripts as described above. Prior to model training, the ‘caret’[39] (version 6.0-64) createDataPartition function was used to split the full cross-cancer and PIC-only data sets into a training cohort containing 70% of patients and a testing cohort containing 30% of patients, while conserving a roughly equivalent number of shortest and longest overall survival patients within each partition.

#### LASSO

A LASSO regression model was trained on the full cross-cancer and PIC and non-PIC only training cohorts using the glmnet[40] (version 2.0-2) cv.glmnet function with regression family set to “binomial” and nfolds set at 5. This generated a binomial regression model, which used a lambda penalty optimized using 5-fold cross validation within the training cohort. The optimal lambda penalty was defined as the smallest model with a cross validation mean squared error within one standard deviation from the minimum value.

#### Ridge

A ridge regression model was also trained with the cv.glmnet function with identical parameters as the LASSO model described above, except the alpha parameter was set to 0.

#### Random Forest

A random forest model was trained on the full cross-cancer and PIC only cohorts using the randomForest[41] (version 4.6-12) package. Models were generated with the randomForest function using default settings except mtry was limited to 1000.

#### SVM

A linear support vector machine model was trained on the full cross-cancer and PIC only cohorts using the e1071[42] (version 1.6-7) package. The model was trained using the svm function with kernel set to “linear” and “cross” set to 5. The cost parameter was optimized for each cohort by finding the value that minimized the 5-fold cross validation squared error within the training cohort after trying a series of values ranging from 0. 00001 to 10000.

#### Model Evaluation

Performance was evaluated for each model by test set ROC curve AUC generated by predictions made on the testing cohort using the predict function and the ROCR[43] package (version 1.0-7).

#### Permutation

The significance of model performance in the PIC only cohort for each machine learning approach was assessed by randomly sampling seven cancers, dichotomizing the cohorts, training each model in an identical manner as described above for the PIC only cohort, and comparing ROC AUC curves for each resulting random sample.

#### Full Cohort Performance Assessment

The LASSO model derived from the PIC-only cohort was applied to the full patient cohorts of each individual PIC to assess performance in a non-dichotomized setting. LGG, KIRC, and LUAD had greater than 25 uncensored patients remaining after removing the 18 poorest outcome patients for model training, so for these cancers the model was applied only on patients that were not used to train the original model. Because KIRP, PAAD, MESO, and ACC had a limited number of remaining patients, the PIC LASSO model was applied to the full cohort including patients that were used to train the original model. The top and bottom quartiles of predicted survival were compared using Kaplan-Meier curves as described above.

### Intra-cancer survival modeling

To assess the relative prognostic ability of tumor PI, random gene signatures, and LASSO optimized signatures, each cancer cohort was split into equivalently sized training and test cohorts 100 different ways with the caret createDataPartition function to ensure an equivalent number of censored events between each group. On each training cohort, a multivariate Cox regression model with LASSO for feature selection was trained with the L1 penalty, lambda was selected by 3-fold cross validation by the glmnet package’s cv.glmnet function with family set to “cox” and “nfolds” set to 3. A random number of genes equivalent in size to the LASSO model generated for each training cohort was selected and used to train a Cox regression with the survival package coxph function. Lastly, a Cox regression model was trained on the tumor PIs of each training cohort using the coxph function. The relative performance of the LASSO, Random and PI models was assessed by concordance index in the test cohort using the R survcomp (version 1.18.0) package concordance. index function with method set to “noether”. Relative concordance indices were compared across all 100 splits for each cancer using paired Wilcox tests.

### Drug associations with proliferation index

To correlate sample PI with drug efficacy, EC50 values for 24 drugs and normalized microarray expression data for 486 cancer cell lines was obtained from the Cancer Cell Line Encyclopedia [44]. The specific files used for analysis were “CCLE_NP24.2009_Drug_data_2015.02.24.csv” and “CCLE_Expression_2012009-29. res” (downloaded in June 2016). Proliferation index was calculated in a similar manner as described above by taking the median normalized expression value for each probe set mapping to a gene contained within the proliferation index. To measure impact of drug treatment on PI, expression profile data of MCF7 cells treated with 1309 drugs and their corresponding vehicle controls was obtained from the Connectivity Map data set [45]. The “rankMatrix.txt” file (downloaded in June 2016) was used for downstream analysis. This file consists of a probe set by treatment matrix with each probe set given a ranking (from 1 to the total number of probes – 22,777) corresponding to the magnitude of differential expression of that probe set after treatment with a drug relative to its vehicle control with a ranking of 1 assigned to the highest positive change in expression and 22,777 assigned to the lowest negative change in expression. The relative impact on PI of different treatments was compared by calculating a median ranking for all probe sets mapping to genes used in the calculation of PI for each treatment and subsequently ranking drugs according to the percentage of drugs with a higher PI ranking. Cumulative distribution functions of all PI-probe set rankings for drugs of interest were also compared.

### Breast cancer subtyping

Subtype assignments for patients in the BRCA cohort were obtained from a previous TCGA analysis of breast cancer [46]. The “PAM50 mRNA” column in Supplemental Table 1 was used for those patients who met our criteria for analysis. Principle component analysis was performed using the prcomp function on the BRCA cohort on all variance stabilized transcript data.

### SNV-point mutation analysis

Somatic mutations were obtained from Kandoth, et al.[47] for 12 TCGA ‘Pan-Cancer’ datasets. We found 2,336 patients overlapped from 9 cancers with the TCGA gene expression dataset and obtained somatic mutations for those patients from Kandoth, et al.’s Supplementary Table 2 where the authors used common, stringent filters to ‘ensure high quality mutation calls’ across those samples. Correlations between tumor PI and somatic mutation burden were conducted by log normalizing the sum of all mutations identified for each patient and performing Spearman correlations with patient PI both across and within each cancer type. To identify genes with mutation status associated with PI, we performed Wilcoxon rank tests of PI between tumors containing a missense or nonsense mutation and tumors containing synonymous or no mutation for each gene with at least 5 mutations present in each cancer. This analysis was not performed on cancers with less than 100 genes meeting these criteria (n=3). To identify significant cross-cancer trends, we used Fisher’s combined p-value method on each gene mutated at least 5 times in at least 2 cancers.

## Results

### Proliferation index varies across tissues, cancer types and tumor pathology

RNA-seq and associated clinical annotation data were compiled for 6,581 patients across 19 cancers. To be included in this study, clinical and RNA-seq data for a given cancer must have been available for at least 50 patients and at least 25 patients must have died from the disease to provide uncensored survival information. Examination of the PI within and across tumor types revealed a continuum of index values within each cancer and notable differences between cancers (Figure 1A). A similar analysis of healthy GTEx tissues revealed low PI values in post-mitotic tissues such as skeletal muscle and brain tissue and higher values in EBV-transformed lymphocytes or tissues with high rates of cell turnover such as esophageal mucosa, vaginal epithelium and skin (Supplemental Figure 1). For every cancer with adjacent normal tissue available from TCGA (n=12), the PI was higher in tumor tissue compared to adjacent normal tissue (Wilcoxon, p<0.05). This was also true when comparing tumor tissue collected by TCGA to normal tissue from the same organs and collected by the GTEx consortium (n=9), demonstrating tumorigenesis is accompanied by a characteristic increase in proliferation-related gene expression (Figure 1B).

**Figure 1.**
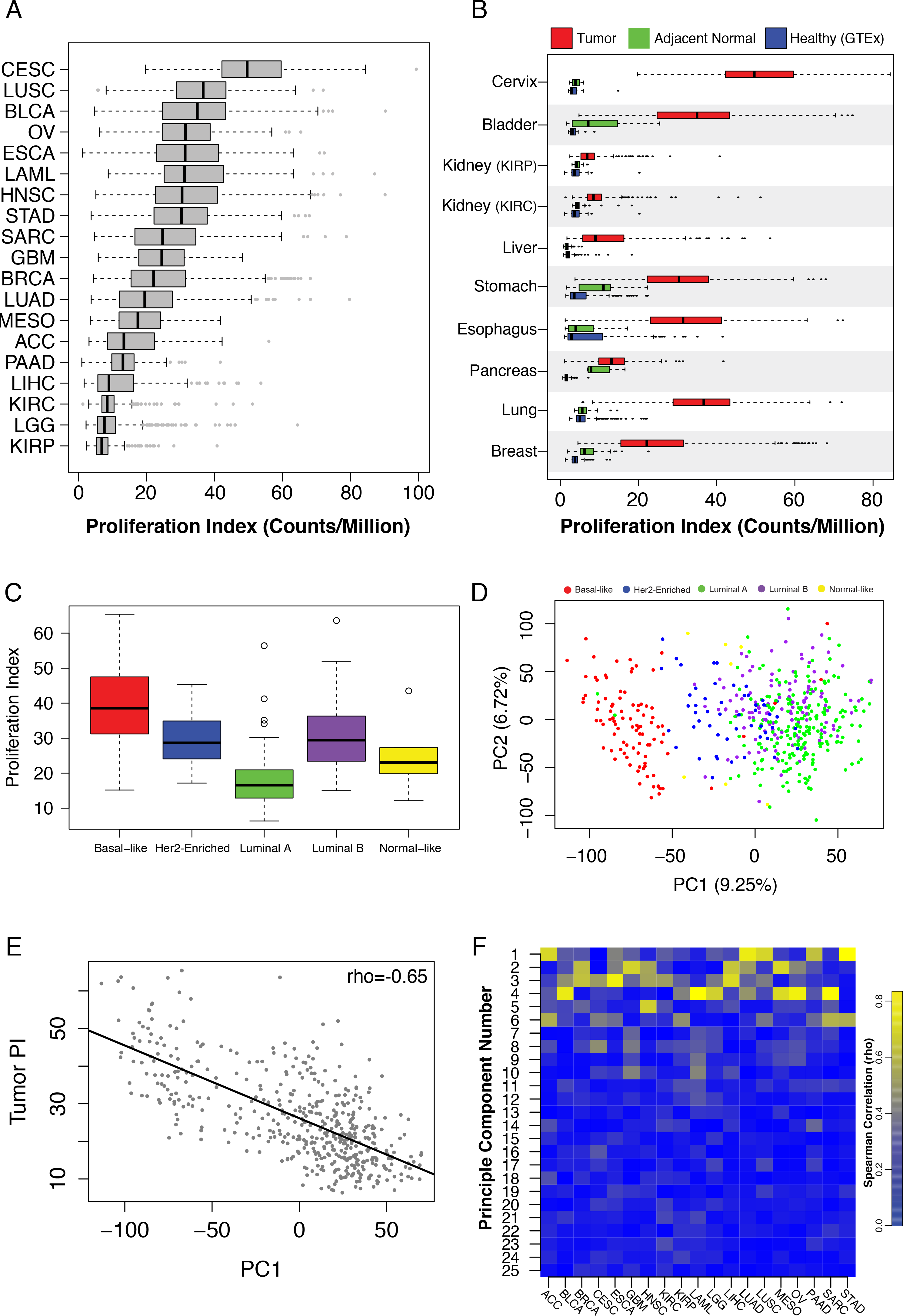
(a) Tumor proliferation index distributions across TCGA cancers. (b) Proliferation index values in healthy GTEx samples (blue), TCGA tumor-adjacent normal tissue (red) and TCGA tumor tissue (green). (c) Tumor proliferation index values across breast cancer PAM50 subtypes. (d) PCA of TCGA breast cancer samples stratifies tumors based on PAM50 subtypes. (e) The first principle component of the TCGA breast cancer data set correlates with tumor proliferation index. (f) Heatmap of principle component-tumor proliferation index correlations across cancers.

Because it was the largest dataset and to follow up the Venet et al. study, we focused additional analysis on breast cancer. Within breast cancer subtypes, PI values were highest among aggressive basal-like tumors and lowest among the less aggressive luminal A and normal-like subtypes (Figure 1C). Principle component analysis (PCA) of all gene expression values in breast cancer confirmed the first principle component (PC1) stratified subtypes (Figure 1D). Interestingly, PC1 was also strongly correlated with tumor PI (rho=0.56) indicating a large proportion of variance within breast cancer gene expression, including subtype delineations, are strongly associated with proliferation (Figure 1E). Moreover, examining PI across all cancers revealed strong correlations with early principle components in a majority of cancers, supporting previous observations that a large portion of variance across tumor transcriptomes is correlated with relative proliferation rates (Figure 1F). However, tumor PI was associated with pathologically assessed tumor stage, nodal invasion, and metastasis in only a subset of tumors analyzed, suggesting the importance of proliferation in tumor progression may vary considerably across cancers (Figure 2A-C). PI values are plotted across each pathological grading characteristic for clear cell renal carcinoma (KIRC), a representative cancer for which PI is significantly associated with pathological stage, and stomach adenocarcinoma (STAD), a representative cancer for which PI is not associated with pathological stage (2D-F).

**Figure 2.**
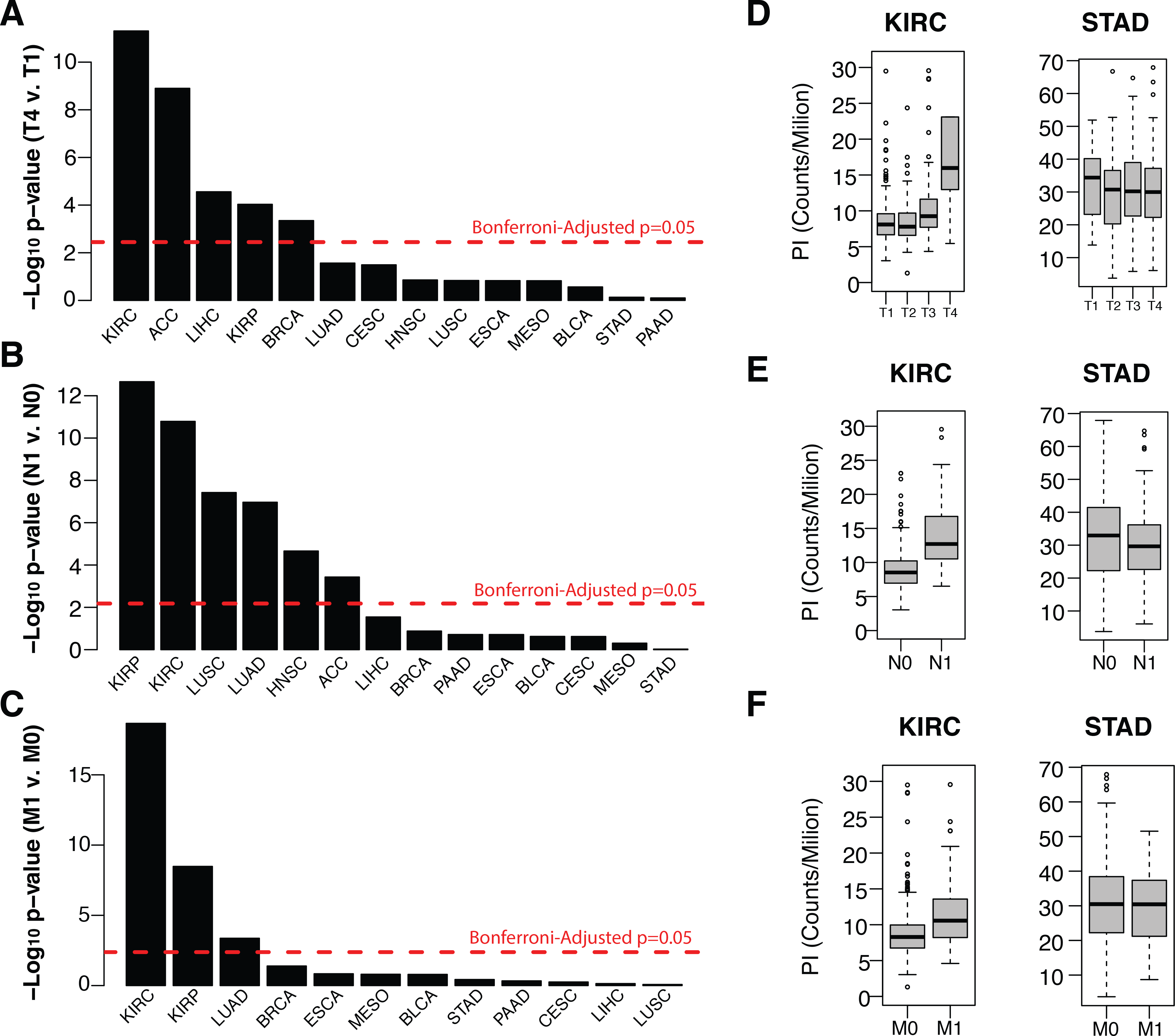
(a-c) Wilcox test negative log p-values of tumor proliferation comparisons between (a) tumor T stages 1 and 4, (b) tumor N stages 0 and 1 (nodal invasion), and tumor M stages 0 and 1 (metastasis)(c). (d-f) Distribution of tumor proliferation index across tumor T (d), N (e) and M stages for TCGA renal cell carcinoma and stomach adenocarcinoma.

### Cell proliferation is associated with overall survival in a subset of cancers

Next we assessed the relationship between tumor PI and patient survival. Cox proportional hazards models and Kaplan-Meier curve analysis revealed tumor PI was significantly associated with survival in a subset of cancers similar to those implicated in Figure 2 above (Figure 3A, Supplemental Figure 2). Strikingly, the association of a cancer’s PI with survival appears to be inversely correlated with its median PI level relative to other cancer types (Figure 3B). This may indicate that in cancers with the highest proliferation rates, other tumor characteristics dictate patient survival. We tested this hypothesis by performing Cox proportional hazards regression on all transcripts in each cancer. Pathway analysis of transcripts significantly associated with survival confirmed an enrichment for proliferation-related gene ontology (GO) terms such as cell cycle, DNA replication, and cell division in cancers whose proliferation rate was associated with survival whereas other cancers showed a relative paucity of proliferation related enrichment and favored cell metabolism, transport, ROS response, angiogenesis and immune related terms (Supplemental Table 3, Supplemental Figure 3).

**Figure 3.**
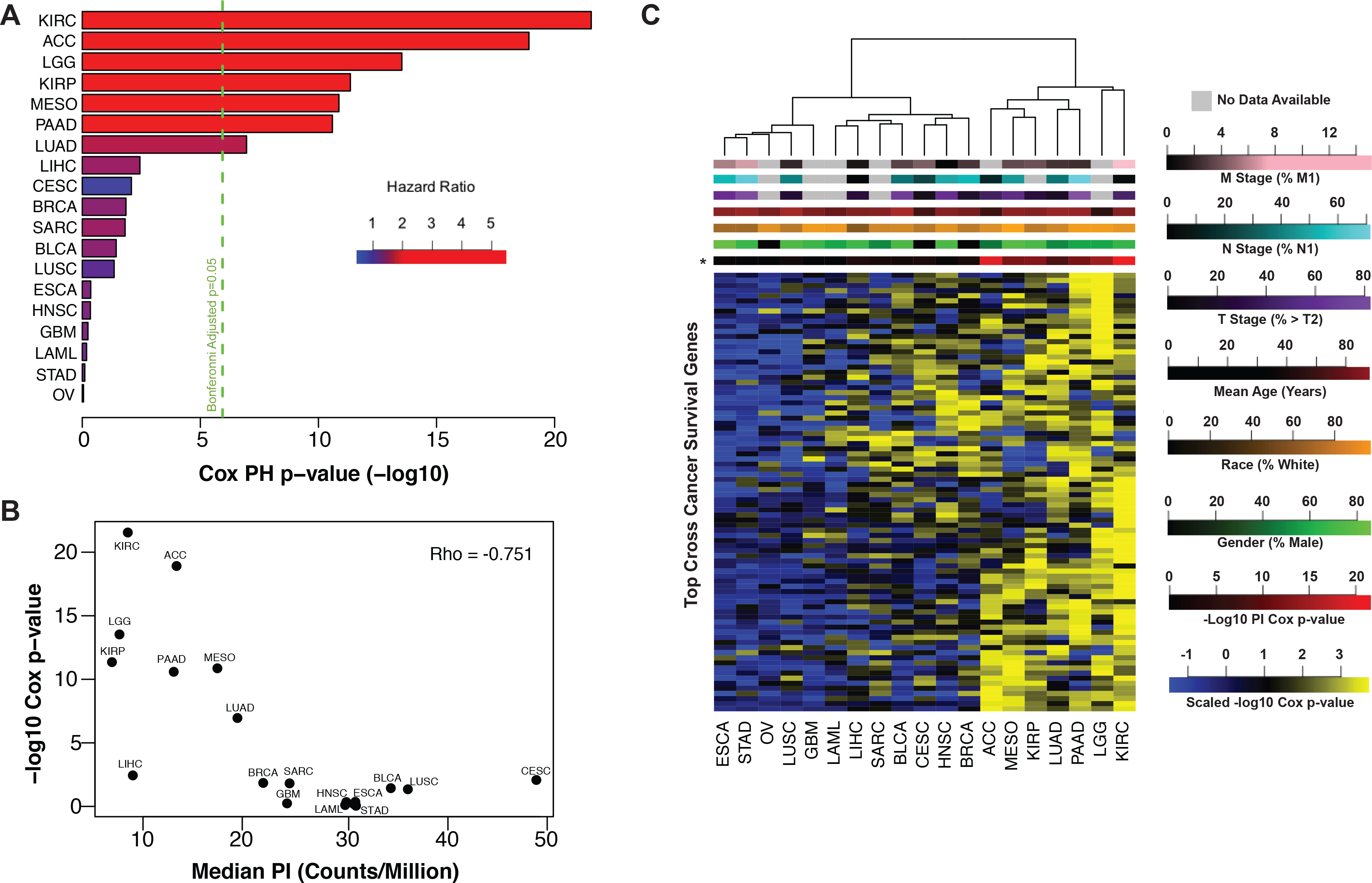
(a) Tumor proliferation index Cox regression negative log p-values plotted by cancer with the first seven cancers showing significant association with patient outcome. (b) Tumor proliferation index Cox regression negative log p-values are anti-correlated with median tumor proliferation index. (c) Heatmap of negative log Cox regression p-values of genes significant (p<0.05, n=84) in at least 9 of 19 cancers identifies PICs (right).

No transcripts were associated with survival in all cancers, however 84 transcripts associated with survival (Cox p-value < 0.05) in at least 9 out of 19 cancers. Pathway analysis on these transcripts revealed enrichment for proliferation-related processes including mitosis, cell and nuclear division, and spindle formation (Supplemental Table 4). Clustering cancers by their respective Cox regression p-values for each of these 84 transcripts revealed two distinct clusters (Figure 3C). The first cluster, representing 12/19 cancers is relatively depleted for low p-values indicating that survival patterns are relatively unique to each of these cancer types. The second cluster, consisting of the remaining 7 cancers, shows a much stronger enrichment for low p-values indicating a common, proliferation-related, survival phenotype. The second cluster of cancers, from here on referred to as proliferation-informative cancers (PICs), is identical to the subset of cancers for which the tumor PI was significantly associated with survival and is not enriched for any clinical or demographic parameter. Relaxing the threshold for the number of significant cancers required for transcript inclusion did not significantly alter this clustering pattern (Supplemental Figure 4). Clustering individual patients based on the expression levels of the top 250 most variable transcripts across all cancers reveals that patients with the same cancer type tend to cluster together and indicates that PIC clustering is not driven by underlying baseline tissue expression patterns (Supplemental Figure 5).

To further investigate cross-cancer survival patterns, we selected an equivalent number of the shortest surviving and longest surviving patients from each cancer and randomly partitioned all samples into training and testing cohorts for prognostic model development and evaluation (Figure 4A). A multivariate Cox regression model with L1-penalized log partial likelihood (LASSO) for feature selection had relatively poor performance (AUC=0.651) when trained on the full set of cancers, however when limited to just PICs, performance improved dramatically (AUC=0.856). This again demonstrates PICs share a common survival phenotype (Figure 4B, Supplemental Table 5). To assess the uniqueness of PICs’ model performance, we randomly selected 1000 sets of 7 cancers for model training and none demonstrated the performance achieved by the PIC-only model (Figure 4C). In fact, model performance across our permutations was strongly correlated with the number of PICs incorporated into each model (Figure 4D). This trend was also observed using a variety of other predictive modeling approaches (Supplemental Figure 6). To assess whether our PIC model could perform well as a continuous metric of survival outside of our pre-dichotomized cohort, we applied it to the full patient cohorts for each PIC. In all PICs, model prediction values were successful in prognostically stratifying patients (Supplemental Figure 7).

**Figure 4.**
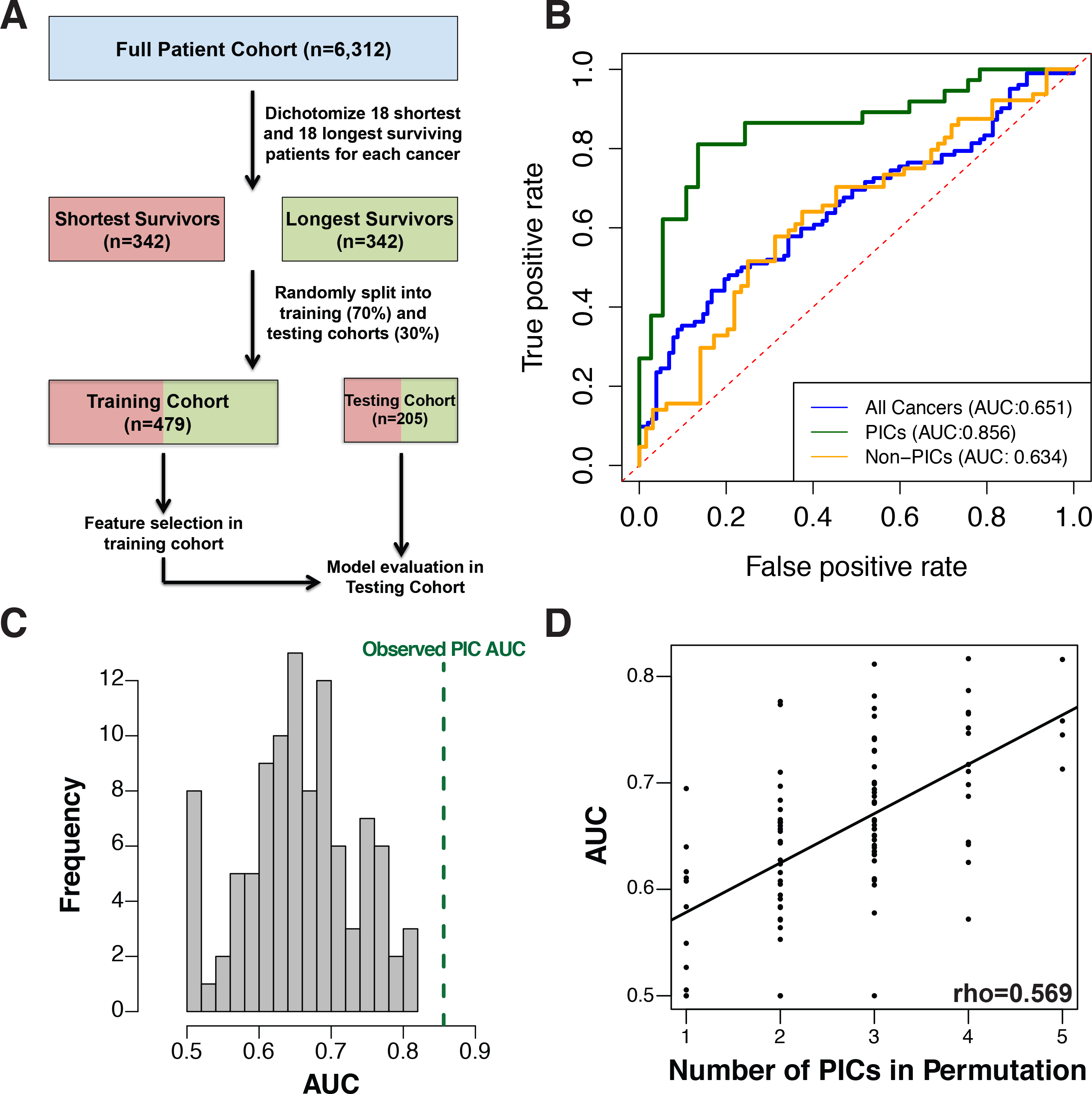
(a) Workflow for cross-cancer survival model generation. (b) ROC curve for multivariate Cox regression with LASSO for variable selection on all 19 cancers (blue), PICs only (green) and non-PICs only (orange). (c) Histogram showing the distribution of ROC curve AUC values for survival models generated on 100 randomly sampled sets of cancers equivalent in number to the PICs. (d) The ROC curve AUC values are directly proportional to the number of PICs included in random sample sets.

We next examined intra-cancer survival patterns by comparing the relative prognostic performance of patient PI, LASSO optimized transcript models, and randomly selected transcript models (Figure 5A). In all cancers except for mesothelioma, optimized transcript models had the best prognostic performance. However in PICs, patient PI and random transcript models were more highly associated with survival than non-PICs (Figure 5B-C, Supplemental Table 6). To facilitate PI exploration, we have developed an R package (https://github.com/blasseigne/ProliferativeIndex), ‘ProliferativeIndex’, which calculates and analyzes PI across a user’s dataset and compares a user’s model with PI.

**Figure 5.**
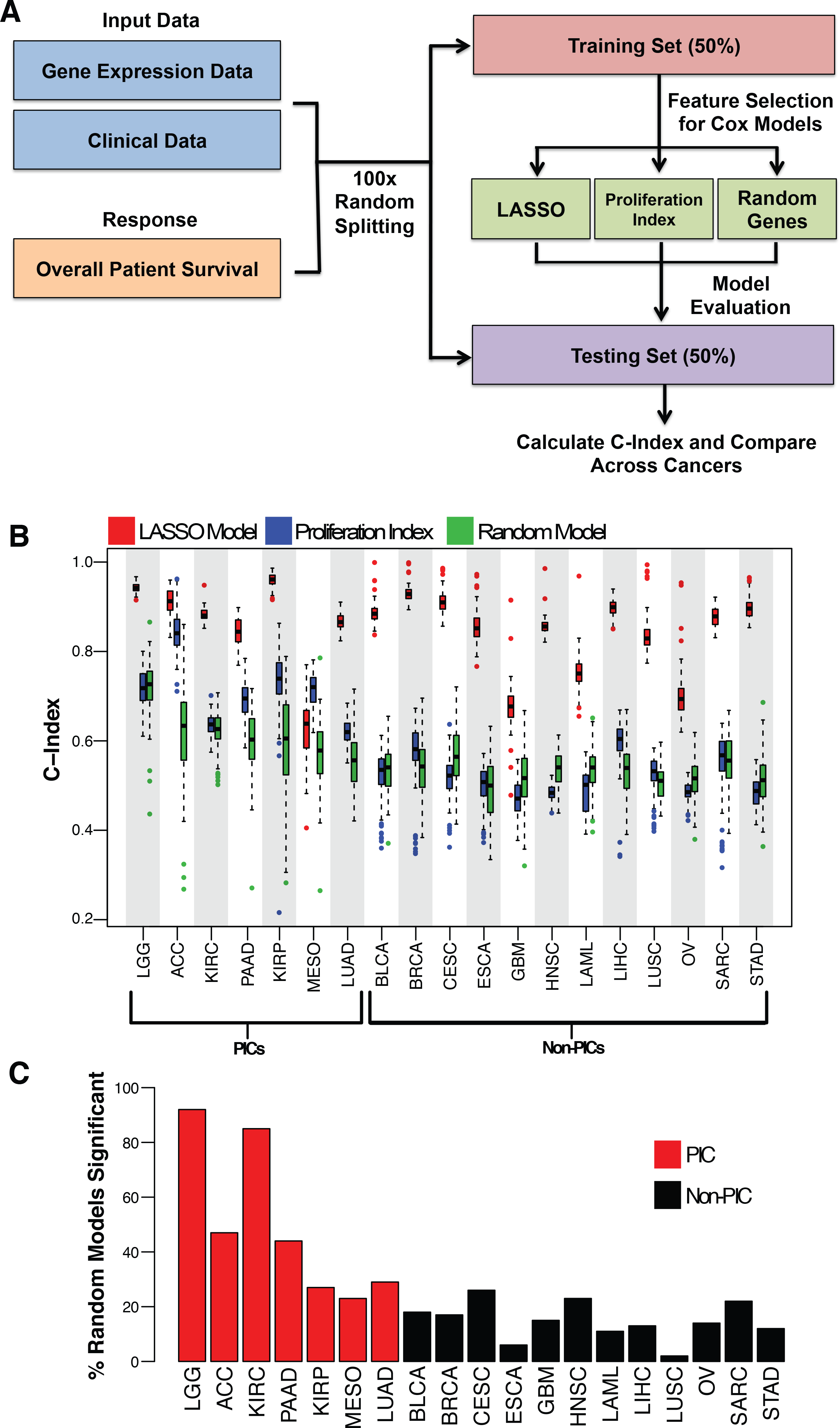
(a) Workflow for generating intra-cancer survival models. (b) Concordance index values for LASSO (red), proliferation index (blue), and random gene (green) models across all TCGA cancers. (c) Percentage of significant random gene survival models based on concordance index across all TCGA cancers.

### Proliferation Index Corresponds to Drug Sensitivity

Because several widely used chemotherapies target proliferation-associated processes, we hypothesized that sensitivity to several drugs may be strongly correlated with PI. To test this hypothesis we obtained drug sensitivity (EC50) and expression information from the Cancer Cell Line Encyclopedia [44]. As suspected, sensitivity to two inhibitors of topoisomerase, an enzyme that controls the unwinding of DNA during replication and transcription, as well as Paclitaxel, a drug that disrupts microtubule function during mitosis, was significantly correlated to cell PI (Bonferroni-adjusted p-value < 0.05, Figure 6A) [48,49]. The EC50 of a histone deacetylase inhibitor, Panobinostat, was also significantly inversely correlated with PI (rho = −0.25, p-value 6.83×10^−7^) and no drug EC50 values were significantly positively correlated with PI. We next investigated whether HDAC and topoisomerase inhibitors were capable of inhibiting proliferation at the transcription level using treatment-induced differential expression data provided for over 1000 compounds by the Connectivity Map (CMap) [45]. We focused on MCF7, a breast cancer cell line for which the most treatment data was available. As a positive control, we examined the effect of estradiol (a ligand previously reported to increase the proliferation rate of estrogen receptor positive MCF7 cells [50]) on PI. We confirmed that nearly all estradiol treatment replicates and concentrations resulted in an increase in PI and found that median PI probe set rank was in the top 20% of all treatments investigated. Conversely, treatment with both HDAC inhibitors present in the CMap database (vorinostat and trichostatin A1) showed the opposite effect with median PI probe set ranking in the bottom 10^th^ percentile of all drugs tested (Figure 6B). Additionally, the cumulative distribution function of PI probe rankings for HDAC inhibitors was significantly lower than estradiol, indicating HDAC-inhibitors down-regulate expression proliferation associated gene expression (Figure 6C). Two topoisomerase inhibitors, etoposide and irinotecan, also fell in the bottom 10^th^ percentile, and etoposide was the most potent inhibitor of proliferation-associated expression of all CMap drugs (Supplemental Table 7). However, several other topoisomerase inhibitors did not follow this trend, suggesting only a subset of drugs in this class are capable of inhibiting proliferation at the transcription level (Supplemental Figure 8). Paclitaxel also did not confer a decrease in PI. It was in the top half of chemo-induced PI increases, a trend consistent with drug activity at the level of microtubule stability rather than transcription. It is possible there is a slight feedback response involving an increase in proliferation-associated expression with paclitaxel.

**Figure 6.**
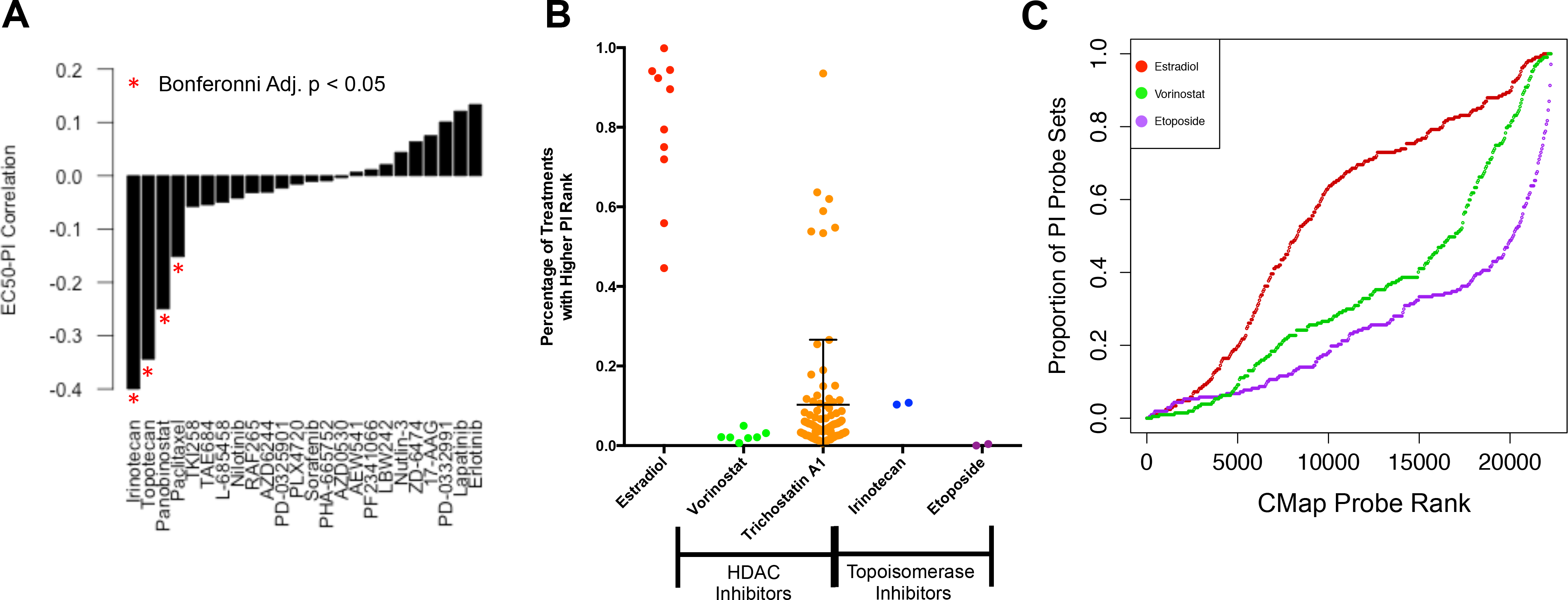
(a) Spearman correlation of EC50 values and PI for 24 compounds across 486 cancer cell lines in the Cancer Cell Line Encyclopedia. Red stars indicate compounds whose correlation is significant (p-value < 0.05) after Bonferonni correction. (b) Treatment induced changes in PI for compounds of interest. CMap rank corresponds to the relative magnitude of differential expression of a probset after treatment with a compound of interest compared to a vehicle control with high rank corresponding to up-regulated genes and low ranks corresponding to down-regulated genes. PI rank was defined as the median rank of probe sets corresponding to PI genes. PI ranks are plotted as percentage of drugs (n=1309) possessing a higher PI rank or decrease in PI after treatment. Mean and standard deviation bars are plotted for Trichostatin A1 (c) Cumulative distribution plots for CMap rankings of probe sets corresponding to PI genes.

### Proliferation and somatic mutation burden

Increased rates of cell division, particularly in cancer cells whose repair mechanisms are diminished, might be expected to correlate with mutation burden. We assessed the relationship between tumor proliferation (PI) and somatic mutation burden in tumor exomes generated by TCGA and previously analyzed by Kandoth et al.[47]. We found a strong correlation between tumor PI and the number of somatic mutations both across and within each cancer (Supplemental Table 8). Notably, total mutation burden and PI were most strongly associated in breast cancer (rho=0.45, Figure 7A). Correlations were also strong within each breast cancer subtype (rho>0.3) except for Her2-Enriched tumors (rho<0.025). We next examined single nucleotide variations (SNVs) most strongly associated with proliferation and found three well-established cancer driver genes enriched for proliferation association mutations (*TP53*, *RB1*, and *PI3K*) consistently implicated across cancers (FDR<0.1, Supplemental Figure 9 and Supplemental Table 9). Apart from these top driver genes, mutations associated with proliferation were tumor-specific. One particular gene of interest, *Reelin*, was among the top 5 genes in breast cancer ranked by protein altering mutations associated with increased PI values in each subtype (Figure 7B). Breast cancer patients within the basal-like subtype tended to have shorter survival times if their tumors harbored protein altering mutations or were low expressers of *Reelin* compared to patients with tumors expressing *Reelin* at high levels (p=0.08, Figure 7C).

**Figure 7.**
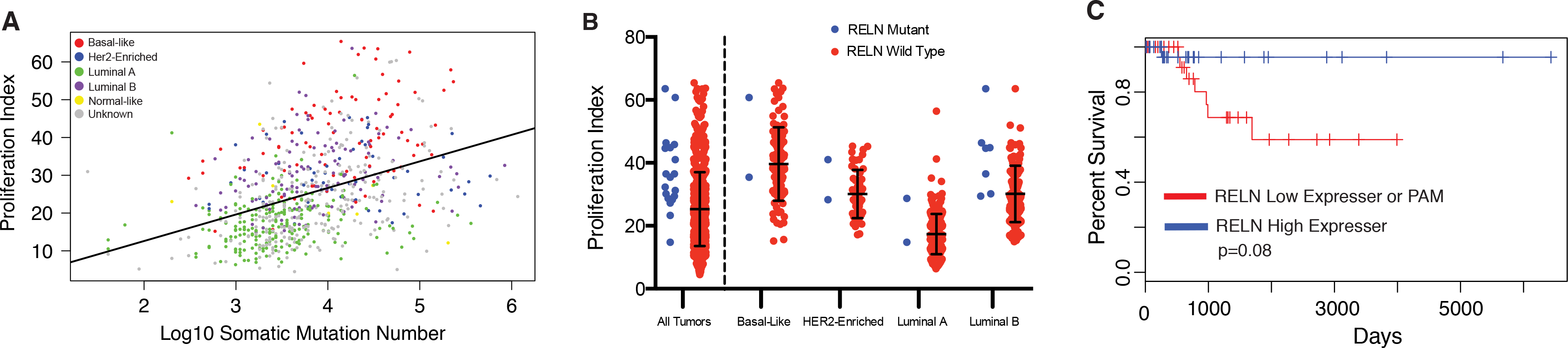
(a) Tumor proliferation index is correlated with TCGA breast cancer somatic mutation burden. (b) TCGA breast tumors containing non-synonymous mutations in *Reelin* have higher proliferation index compared to wild-type. (c) Kaplan-Meier survival plot shows reduced expression or protein altering mutations in RELN are markers of poor prognosis in patients with basal breast cancer.

## Discussion

We have described an RNA-seq based analysis of cell proliferation across 19 cancers in 6,581 patients. We show a high degree of variability in the relative expression of proliferation-associated genes both within and across cancers. Interestingly, cancers with relatively low expression of proliferation-associated genes tended to be those for which PI was strongly associated with pathology-based markers of tumor staging and survival. This suggests that some cancer types may saturate their capacity for proliferation at early stages, allowing other factors such as invasion, immune suppression, and drug transport to dictate prognosis. Proliferation may play a more prominent role in dictating prognosis in cancers that avoid maximal rates of cell division during early tumorigenesis and possess relatively lower absolute levels of proliferation-associated expression. Future studies investigating evolutionary histories of tumors could investigate this phenomenon in more detail as there may be considerable heterogeneity between cancers in the genes important to predicting patient survival and it is possible the most effective targeted therapy would target pathways most relevant to patient outcome. We have demonstrated that PI is significantly correlated with the sensitivity to a subset of drugs *in vitro* and have highlighted several drugs capable of reducing the expression of proliferation-associated transcripts, however future studies are necessary to confirm the relevance of these observations at physiologically constrained doses *in vivo*.

Additionally, we demonstrated survival-associated gene expression patterns were not common across all cancers. However a subset of cancers, PICs, share an overlapping signature enriched for proliferation-associated genes. We were able to develop a common prognostic signature that accurately predicts patient survival across all seven PICs. This survival signature contains several genes previously implicated in cancer prognosis. For example, *CKS2* is a regulatory protein that binds the catalytic subunit of cyclin-dependant kinases and is essential for kinase function in regulating the cell cycle[51,52]. *CRYL1* has been shown to regulate G_2_-M phase transition and expression has been linked to patient prognosis [53]. *DNA2* is a DNA helicase that plays an important role in processing Okazaki fragments during DNA replication and *DNA*2 expression has been correlated to patient survival [54]. *HJURP* is a histone chaperone shown to play a role in the progression of gliomas and breast tumors [55,56]. *SUOX* had the largest absolute coefficient in our model, however its role in cancer progression is less clear. It is a mitochondrial enzyme that catalyzes the conversion of sulfite to sulfate and has been described in one study as a prognostic immunohistochemical marker for hepatocellular carcinoma [57], yet its functional importance in cancer remains unclear. Notably, a significant number of random transcript models generated for PICs were able to significantly predict patient survival indicating that a large portion of the transcriptome is likely correlated with survival in these cancers. Future prognostic modeling within PICs or cross-cancer modeling including PICs should consider the significant role of tumor proliferation-associated expression before interpreting biological mechanisms for prognostic-associated genes. Additionally, newly developed prognostic models in PICs should outperform general transcriptome associations with survival before mechanistic interpretations are made.

Proliferating tumors, which must constantly replicate their genomes, are prone to increased mutation rates, therefore it follows logically that tumor PI is strongly correlated with somatic mutation burden both within and across cancers. This may provide a potential mechanism by which increased proliferation rates associate with poor outcomes as increasing the mutational heterogeneity of a tumor may lead to avenues of escape from targeted drug therapies [58]. Correlation of gene mutation burden with tumor PI revealed three well known tumor suppressor genes (*TP53*, *RB1*, and *PI3K*) to be significantly associated with proliferation across multiple cancers, confirming large bodies of previous work. For example, a large analysis of *TP53* levels in node-negative breast cancer revealed decreases in *TP53* were strongly associated with a concurrent increase in both tumor proliferation and poorer patient outcomes[59]. Moreover, an extensive body of literature exists examining *PI3K*’s ability to upregulate proliferation machinery through downstream activation of the *AKT/mTOR* pathway[24]. Focusing on breast cancer, the largest cancer cohort available, we found one relatively less investigated gene, *Reelin*, among the top PI associated genes. We found that protein-altering mutations in *Reelin* were associated with increased tumor PI in each breast cancer subtype, and that low levels of *Reelin* expression were associated with poor prognosis within the basal subtype. *Reelin* could hold particular interest in breast cancer as its expression in the hippocampus has been shown to be downregulated in response to corticosteroids often given to breast cancer patients in an effort to mitigate the negative effects of chemotherapy. *Reelin* expression has also been shown to be linked to dopamine signaling and may help explain recent successes in targeting dopamine receptor-1 *in vitro* and increased cancer risks in patients receiving antipsychotic dopamine antagonists [60–62]. Although a majority of investigation has been carried out in brain tissue, there may be intriguing roles for *Reelin* in the progression of breast cancer. In sum, this analysis provides a comprehensive characterization of tumor proliferation rates and their association with disease progression and prognosis across cancer types and highlights specific cancers that may be particularly susceptible to improved targeting of this classic cancer hallmark.

## Supplemental Figure Legends

**Supplemental Figure 1.**
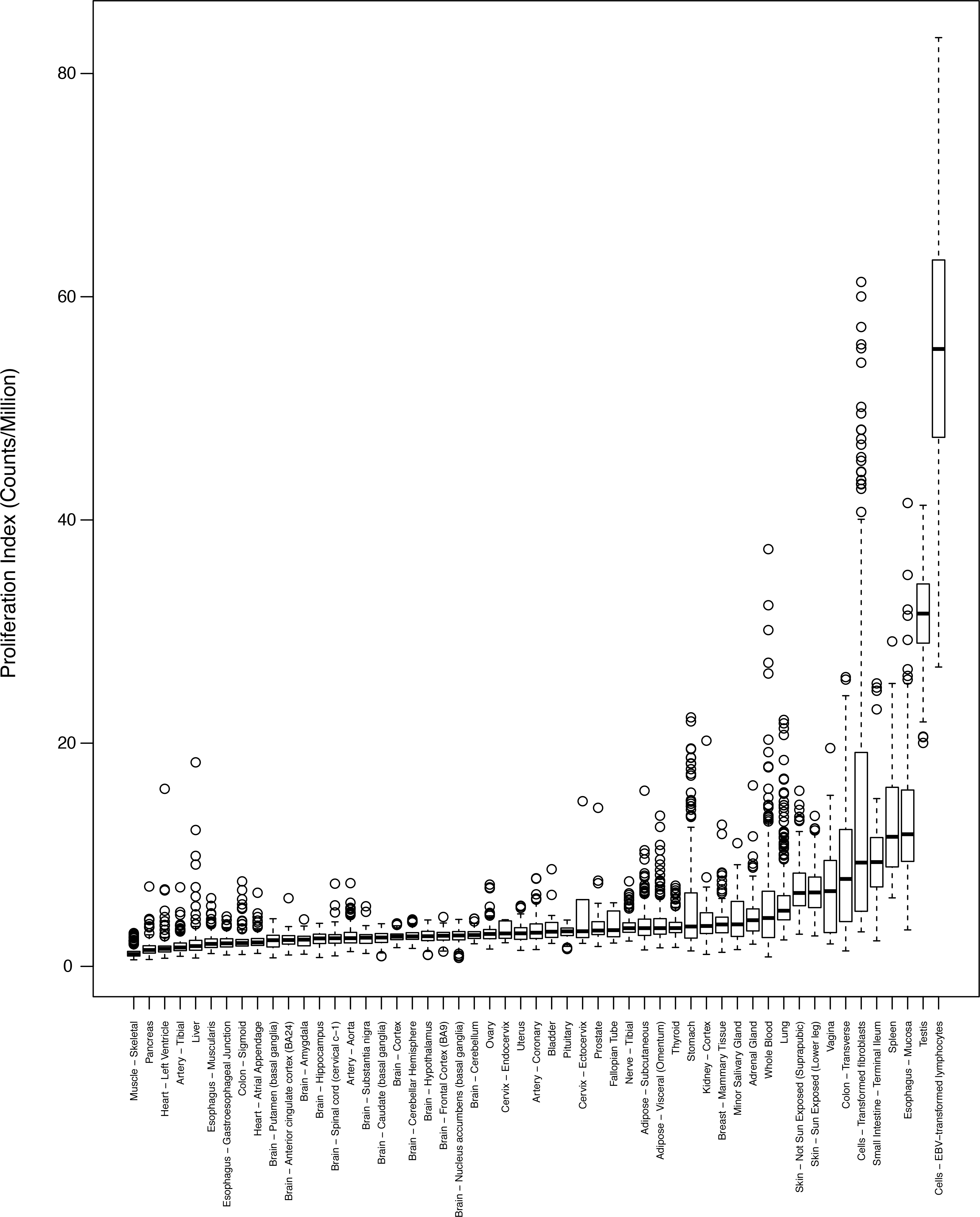
Proliferation index distributions across GTEx healthy tissues.

**Supplemental Figure 2.**
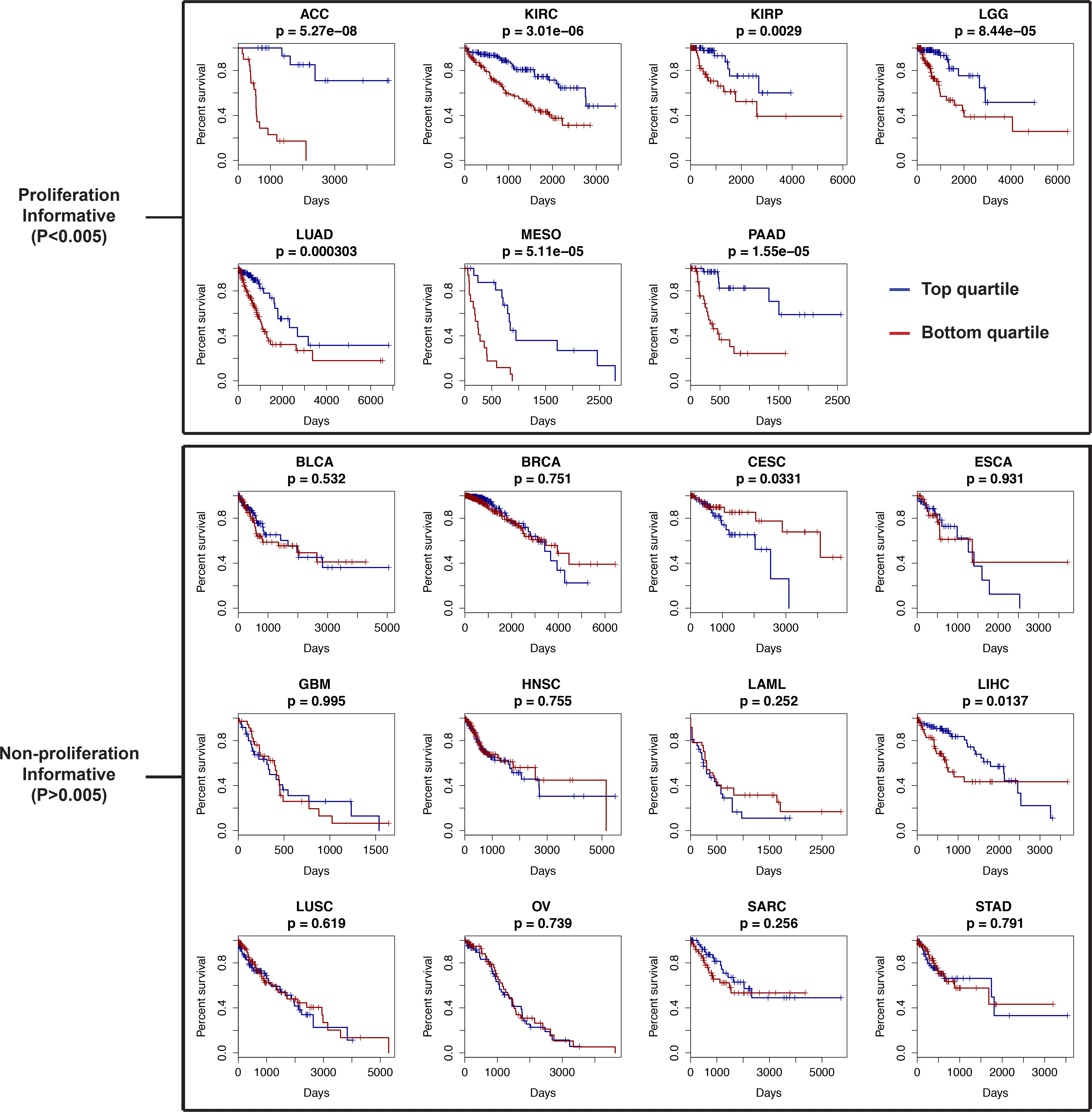
Kaplan-Meier curves for the top and bottom quartiles of tumor proliferation index across TCGA cancers.

**Supplemental Figure 3.**
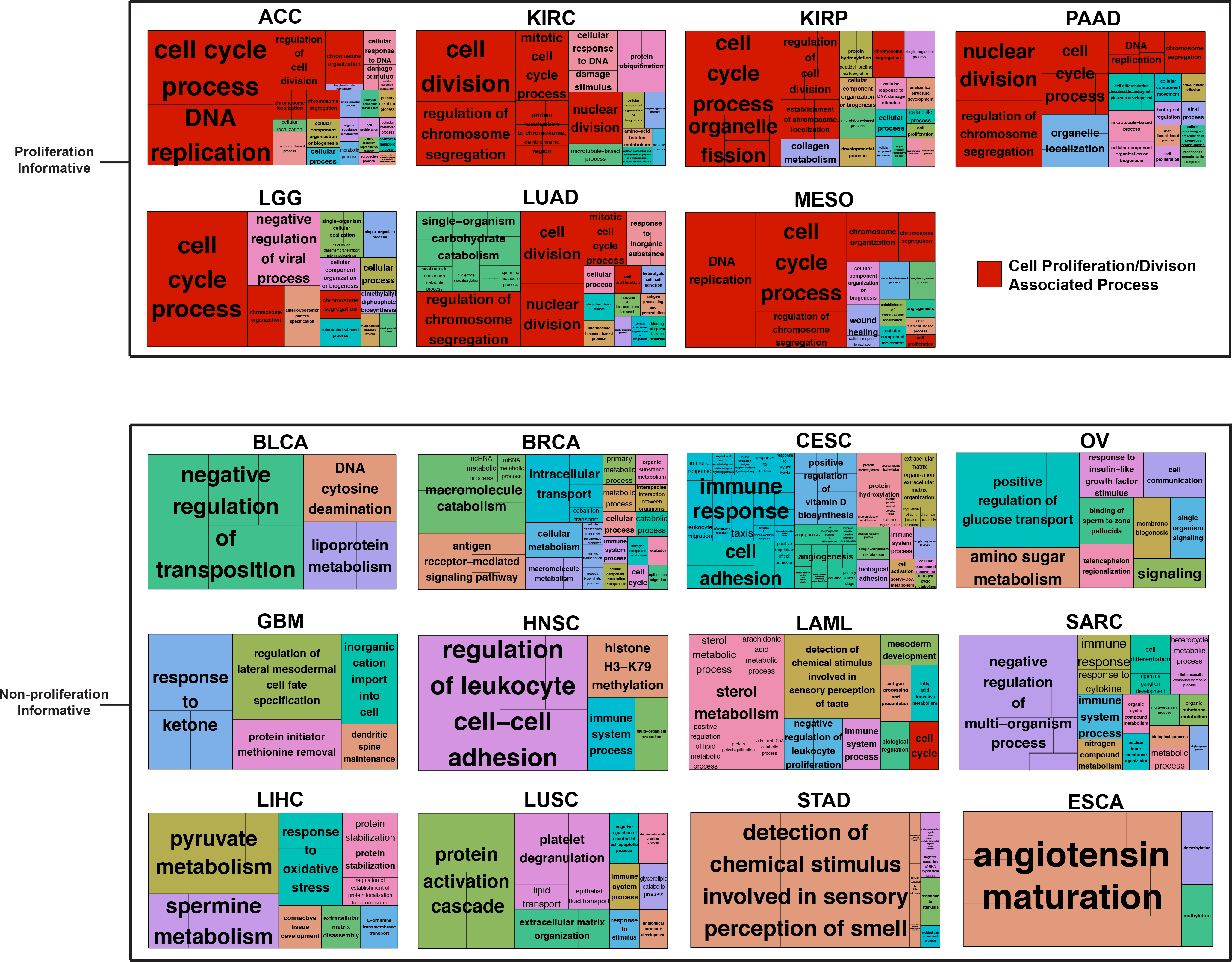
Gene ontology enrichment analysis on survival associated genes in each TCGA cancer. Cell proliferation and division associated modules are colored in red. Module size is proportional to the number of significant genes contained within the module.

**Supplemental Figure 4.**
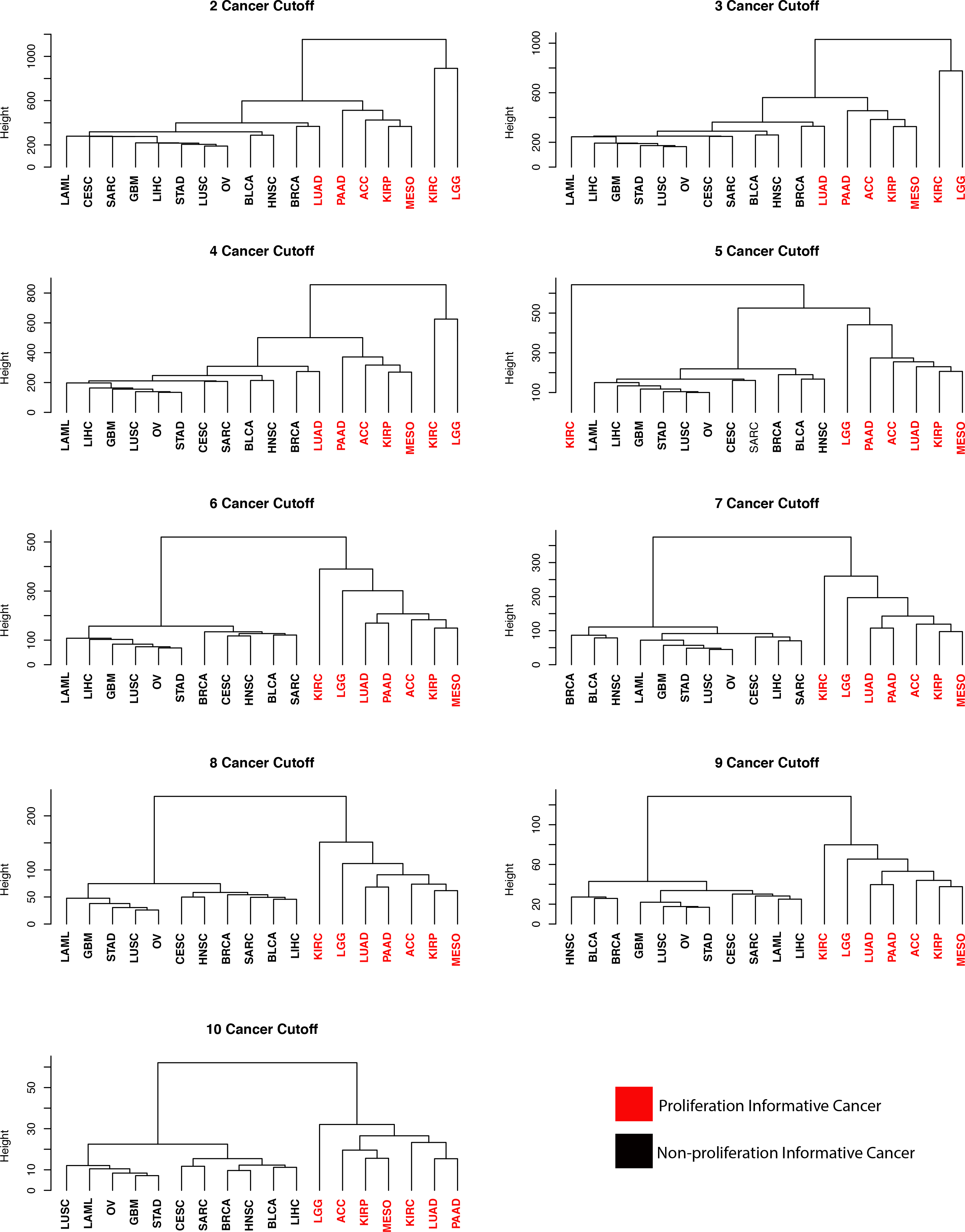
Dendrograms showing cancer clustering based on survival associated p-values using a sliding cutoff for inclusion of genes. Gene inclusion cutoff ranged from requiring genes to be significant (Cox p<0.05) in at least 2 cancers to at least 10 cancers. The PIC clustering pattern in Figure 3 is maintained across cutoffs.

**Supplemental Figure 5.**
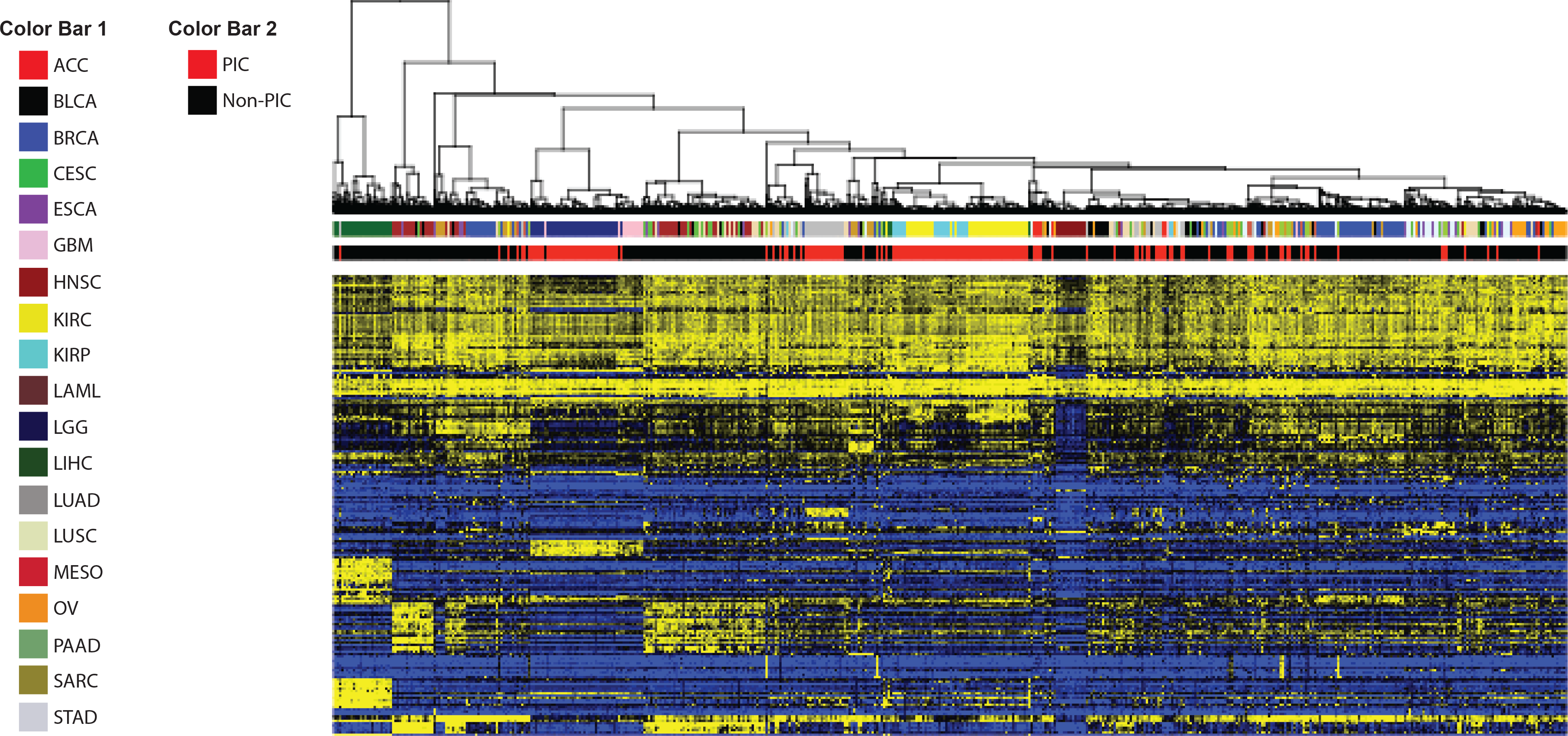
Tumor sample clustering based on expression levels of the top 250 most variable genes across all TCGA samples included in our analysis. Patients with the same cancers tend to cluster together despite PIC status.

**Supplemental Figure 6.**
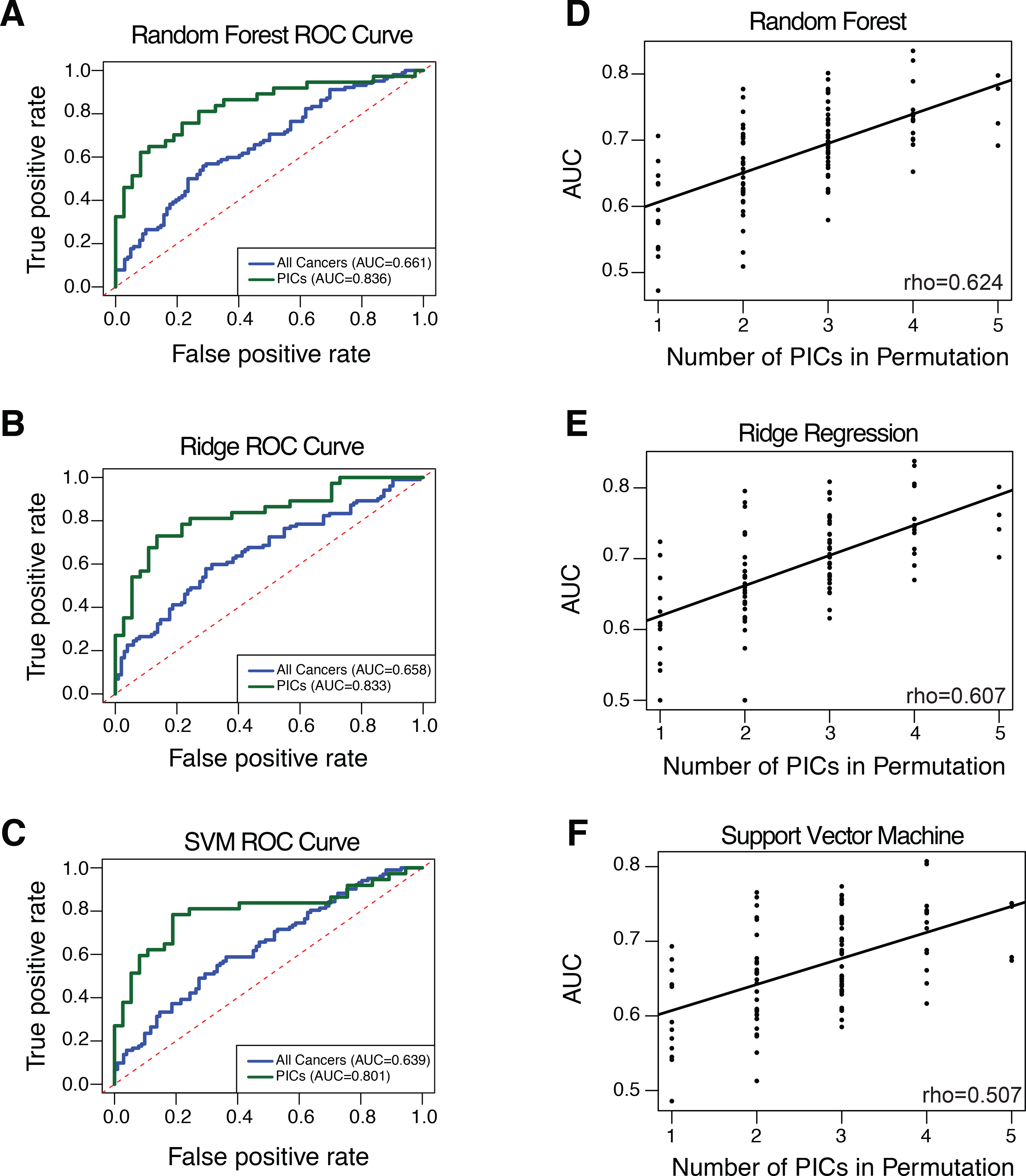
Cross-cancer survival model performance generated on all cancers (blue) and PICs (green) with random forest (a), ridge regression (b), and support vector machines (c). (d-f) ROC curve AUC values from survival models generated on random sets of cancers equivalent in number to PICs are directly correlated with the number of PICs included in the random sample for each survival modeling approach.

**Supplemental Figure 7.**
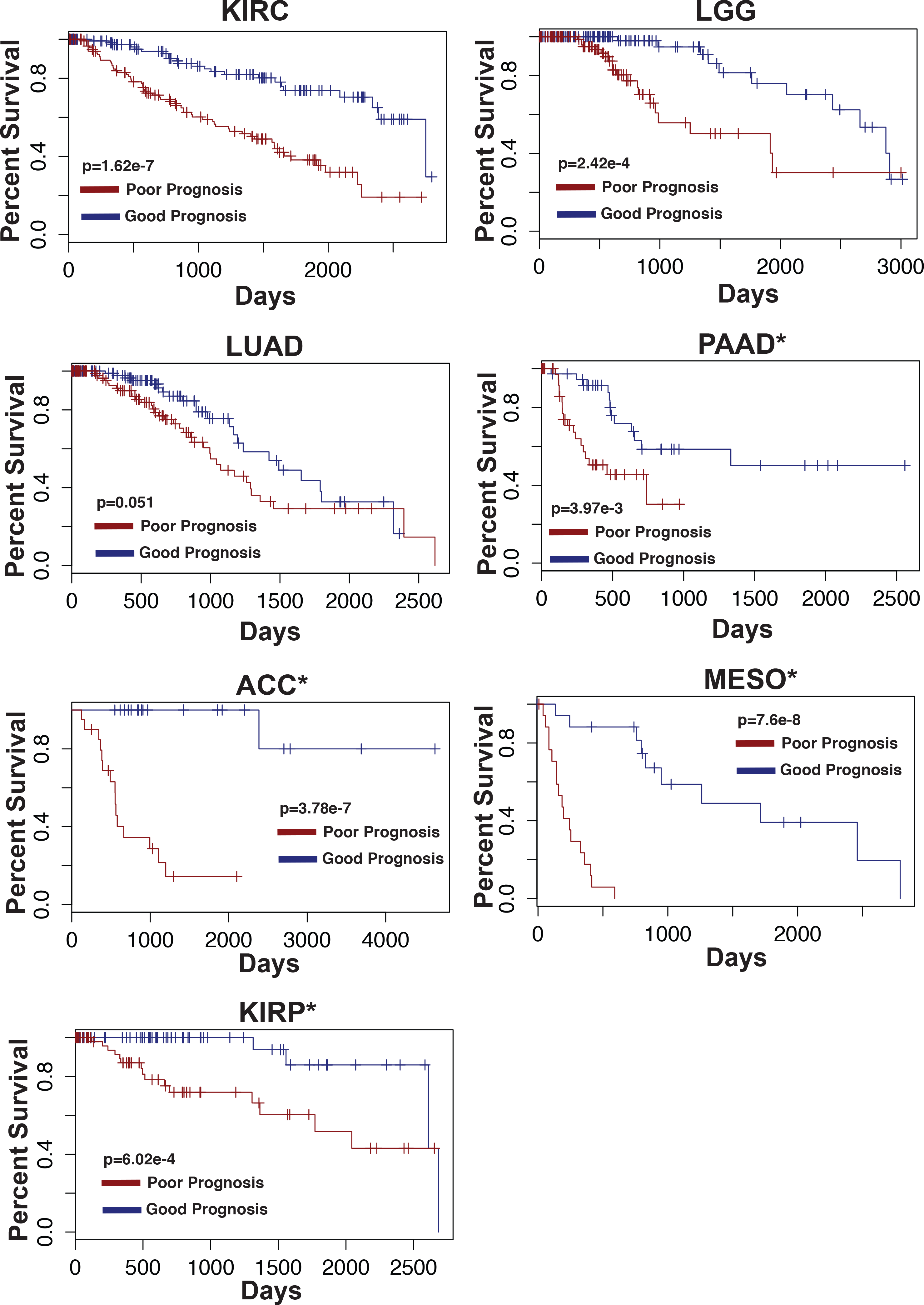
PIC LASSO based cross-cancer survival model performance on the full cohorts for each PIC. Kaplan-Meier curves represent the top and bottom quartiles of patients in terms of predicted prognosis. PAAD, ACC, MESO, and KIRP had an insufficient (<25) number of patients remaining after removing patients used to train the model, thus those Kaplan Meier curves include patients used in the initial training set.

**Supplemental Figure 8.**
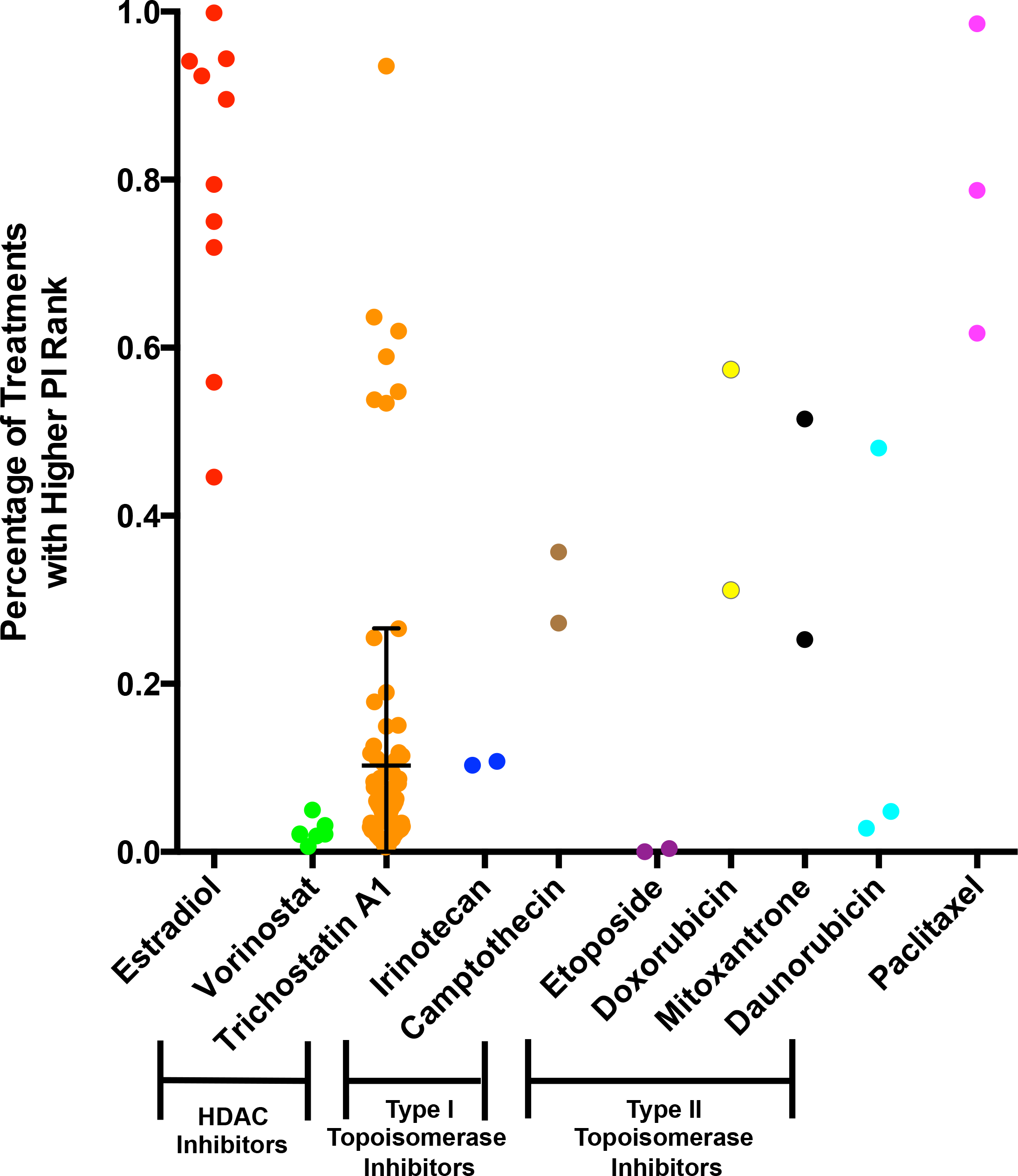
PI ranks, plotted as percentage of drugs (n=1309) possessing a higher PI rank or greater increase in PI after treatment, for all drugs of classes implicated in Figure 6A and estradiol. Mean and standard deviation bars are plotted for Trichostatin A1

**Supplemental Figure 9.**
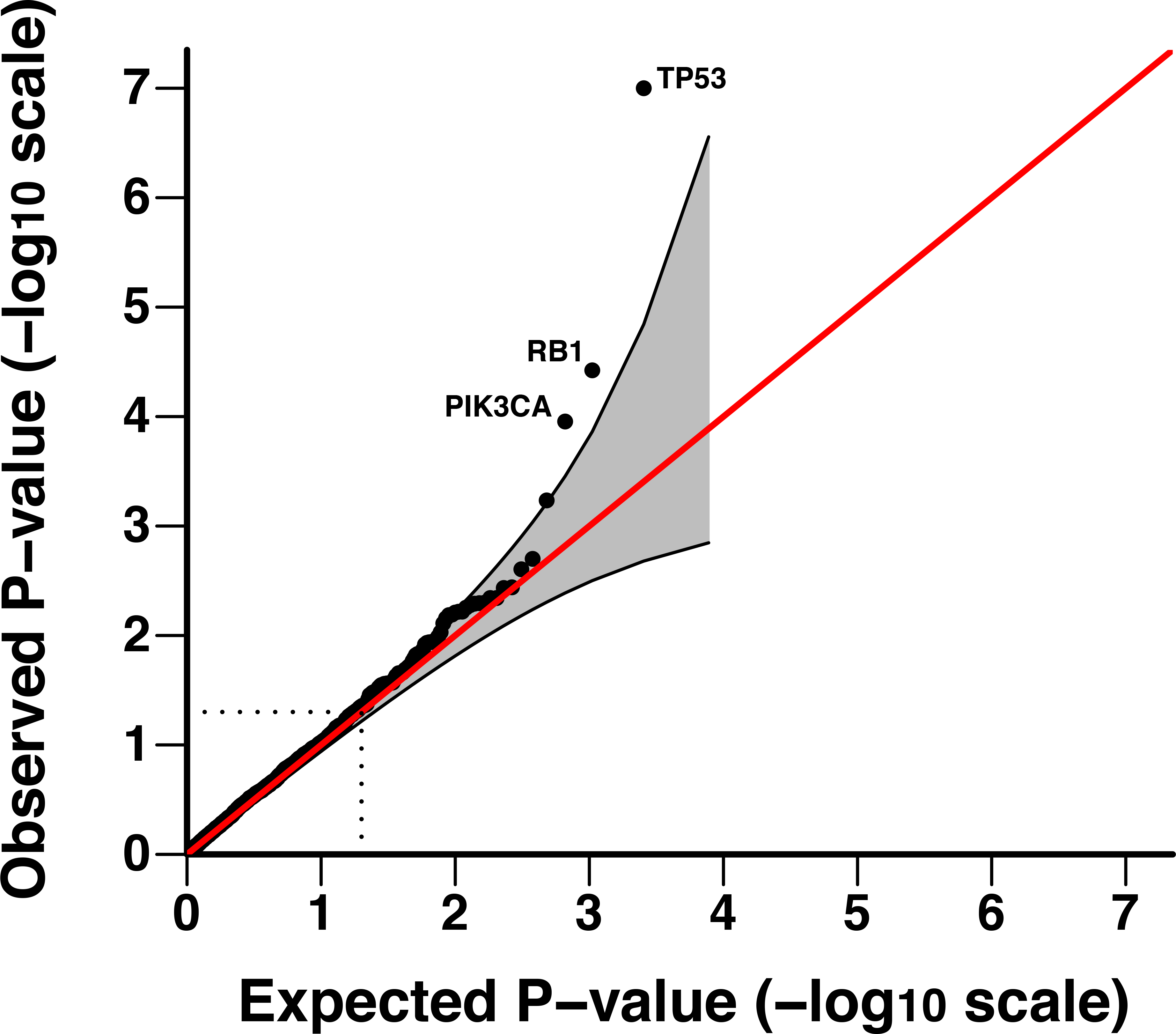
Q-Q plot of p-values derived from gene mutation burden-proliferation index associations.

